# Music for cells? A systematic review of studies investigating the effects of audible sound played through speaker-based systems to cell cultures

**DOI:** 10.1101/2020.11.02.364653

**Authors:** Dongho Kwak, Thomas Combriat, Chencheng Wang, Hanne Scholz, Anne Danielsen, Alexander Refsum Jensenius

## Abstract

There have been several studies investigating whether musical sound can be used as cell stimuli in recent years. We systematically searched publications to get an overview of studies that have used audible sound played through speaker-based systems to induce mechanical perturbation in cell cultures. A total of 12 studies were identified. We focused on the experimental setups, the sound materials used as stimuli, and the outcomes. The stimuli were categorized into simple and complex sounds. The effects were reported as enhanced cell migration, proliferation, colony formation, and differentiation ability. However, there are significant differences in methodologies and cell type-specific outcomes, which made it difficult to find a systematic pattern in the results. We suggest that future experiments should consider using: 1) a more controlled acoustic environment), 2) standardized sound and noise measurement methods, and 3) a more comprehensive range of controlled sound stimuli.

## Introduction

Music for cells. Is there such a thing? The use of (musical) sound to induce a beneficial physiological and psychological effect on biological beings is probably not a novel idea. Already some of the Greek philosophers, such as Pythagoras, recognized the fascinating relationship between musical sound and the human body (1,2). With more advanced technological development, the efforts to attain a deeper understanding of the relationship between sound and body have become more specific, now at a minuscule level in the field of mechanobiology. Mechanobiology has been developing substantially in the last few decades with more than 33,000 publications in the field up until 2016 (3,4). More recently, there have been escalating interests and efforts to understand mechanotransduction through mechanical cues induced by acoustical perturbation on cellular responses (5). Various cell types were investigated including: plant cells (6), micro-organisms (7), auditory cells (5) and other mammal cells *in vitro* (8) or *in vivo* (9). However, the cell types, experimental setups, sound measurement methods, and outcomes are widely dispersed in the literature. In this review, the aim is to get an overview of studies that have used audible sound played through speaker-based systems to induce mechanical perturbation in cell cultures (particularly bacteria and mammalian cells). The paper starts with a background to the mechanobiology of cells and some fundamental physical properties of sound. A review of the literature focuses on the experimental setups, the sound material used (categorized into simple and complex sounds), and the experimental outcomes.

### Background

#### The mechanobiology of cells

Cells *in vivo* are endlessly perturbed by different types of mechanical forces such as tension, compression, and shear stress (10), and it has been suggested that cells are sensitive to physical forces (11). Mechanical signals, such as adhesiveness or extracellular matrix (ECM) stiffness, are also sensed by cells (12). Different types of mechanical cues instigate specific responses in cells where they activate certain genes through distinct signaling pathways (13). For instance, the Hippo pathway has been identified as conserved mechanical transducers, which is critical for driving stem cells fate and regeneration, translocate to the nucleus in cells that are subjected to a stiff surface (12,14). At the same time, its altered activity is involved in aberrant cell mechanics transduction and several diseases (15). Cells sense not only the mechanical signals but also generate signals themselves. It has been reported that bacterial cells can generate sound waves between 8 and 43 kHz (16), yeast cells can generate mechanical vibrations (17), and stem cells have shown unique vibrational signatures to sound stimulation (18).

#### Sound as a mechanobiological stimulus

Sound is essentially vibration. The word *vibration* originates from the Latin word *vibrationem,* which means shaking. When an object is disturbed by mechanical stimulus and starts shaking and vibrating, sound energy is produced (19). This energy is propagated in the form of pressure waves in all directions from the sound source, usually as a longitudinal wave consisting of compressions and rarefactions of matter. In some cases, sounds in solid mediums can travel as a transverse wave, but also as a longitudinal wave. Such a wave consists of up and down motion perpendicular to the direction of the vibration. Sound waves can travel through different mediums, including gas, liquid, and solids, but with different speeds dependent on the medium. Typically, sound travels faster in a liquid or solid medium, for example, water or a steel bar, than in gas medium (20–23).

A medium’s density impacts energy transmission when sound is travelling from one medium to another. This physical property of sound is a factor that needs attention in cellular experiments, in which sound energy must be transmitted from the air into a liquid-based solution, or another possible cell medium. For a progressive plane wave the physical property of a medium can be expressed as the acoustic *impedance*, defined in equation 1.

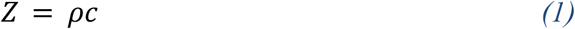

In equation 1, *ρ* is the density (kg/m^3^) of a medium and *c* is the speed of sound (m/s) in the medium. At the interface between two media of mismatched acoustic impedance, part of the incident energy will not be transmitted but will be reflected back to the initial medium. The amount of acoustic energy that will undergo reflection at the interface between a medium of impedance *Z*_*1*_ and a medium of impedance *Z*_*2*_ can be evaluated by the acoustic energy reflection coefficient defined in equation 2.

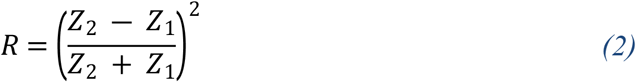

For example, sound pressure waves travelling from the air at 20 °C (*Z*_*1*_≈413 Pa·s/m) will be reflected almost entirely (*R*≈0.9989) at the surface of the water (*Z*_*2*_≈1.5·10^6^ Pa·s/m).

The medium’s thickness also impacts the transmitted energy. If the thickness of the second medium is much smaller than the wavelength of the incident sound, for instance, the amount of transmitted energy will be minimized (19).

#### Manipulatable sound parameters

When exploring the use of sound for experimental set up of cells, there are several properties to consider. Some of the basic parameters of sound include its frequency (what is experienced as *pitch*), amplitude (which corresponds to perceptual *loudness*), and exposure time. Additionally, there are different types of waveforms, from the most basic (sine tones) to more complex, synthetic waveforms (sawtooth, square, triangle), and even more complex, natural waveforms (noise, speech, music). With modern digital signal processing tools, all of these can be created and/or modified systematically in different ways using various types of synthesis techniques (additive, subtractive, frequency modulation, amplitude modulation, etc.) and a broad range of effects and processing techniques in both the time and frequency domains (24).

When it comes to basic descriptions of sounds, the sound frequency (Hz) is represented as the number of cycles or vibrations per second, defined in equation 3.

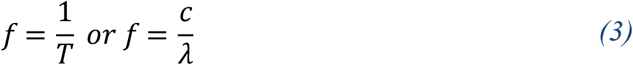

In equation 3, *T* is the period (s) which is the time taken for a vibrating motion to complete one cycle, *λ* is the wavelength (m) which is the distance between two successive matching points in any section of the sound wave cycle. When the period and wavelength increase, the frequency will decrease in a given material, but the speed of sound will remain the same depending on the temperature, in the air, for instance. There is more energy in sound frequencies with a higher amplitude than in sound frequencies with a lower amplitude. Amplitude is the difference between maximum and minimum values of the pressure with respect to the equilibrium pressure, and has a direct influence on the sound pressure level (SPL; dB SPL) (25).

In the following, we will differentiate between “simple” and “complex” sound stimuli. Simple waves, such as a sine tone, are constant and repetitive, and can be produced by “any vibrating body executing simple harmonic motion”. The Fourier theorem tells that any complex sound can be broken down to a given number of sinusoidal components (fundamental and its harmonics) when the signal is periodic. Similarly, any periodic sound can be constructed using a set of sine tones using the Fourier transform. There are also other types such as sawtooth and triangle waves that can be constructed using an infinite set of sine tones (26). Complex waveforms, such as music and speech, are usually not repetitive (21).

## Review method

In this review study, an exhaustive literature search was performed using the University of Oslo Library’s Oria search engine, PubMed, and Google Scholar. The keywords included in the search were: “sound”, “music”, “acoustic stimulation”, and “cell culture”. The combinations of words used for the search were: “sound stimulation on cell culture”, “music stimulation on cell culture” and “acoustic stimulation on cell culture”. Publications were collected between 1 July 2019 and 20 June 2020.

As illustrated in Fig 1, a total of 95 records were identified through the database searching. Of these, 7 records were removed as duplicates, 74 records were investigated but considered not relevant and hence removed. This included studies which were not related to cell studies *in vitro*, or not related to audible sound stimulation, or publications not in English. The full texts of 2 records were unobtainable and were not considered either. This left us with 12 studies that were included for further qualitative consideration (Table 1).

**Fig 1.**
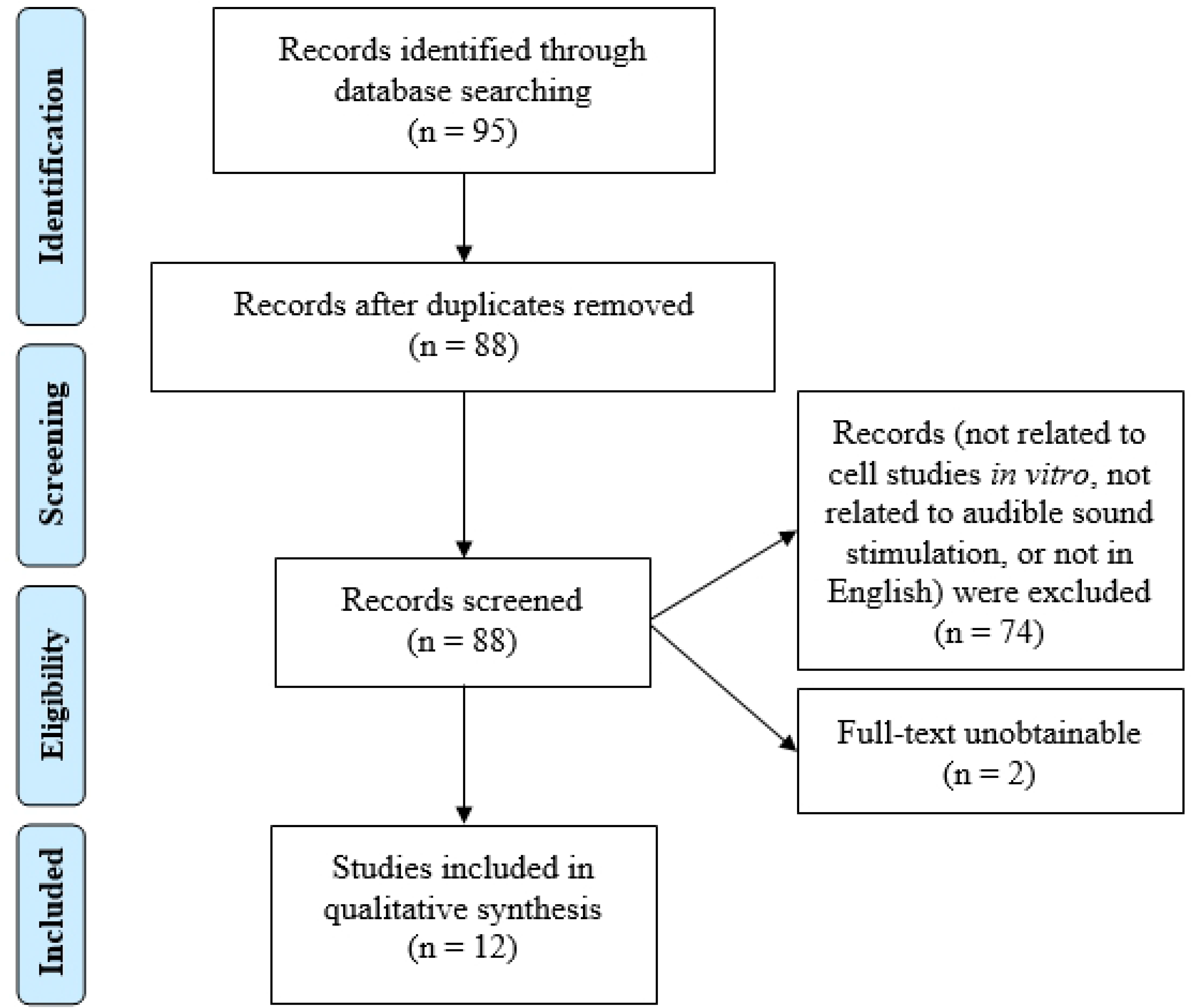
PRISMA flow diagram.

**Table 1.**
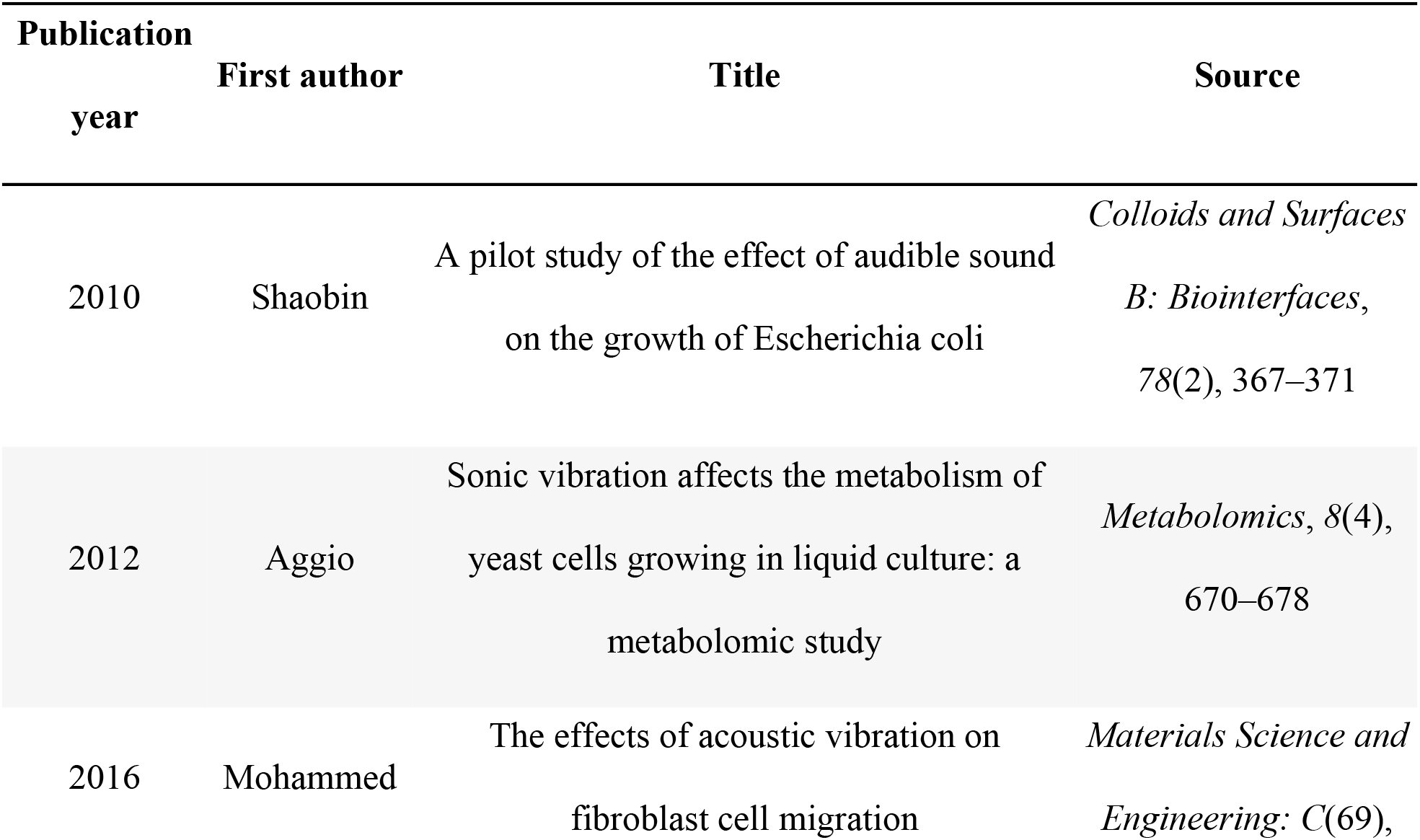
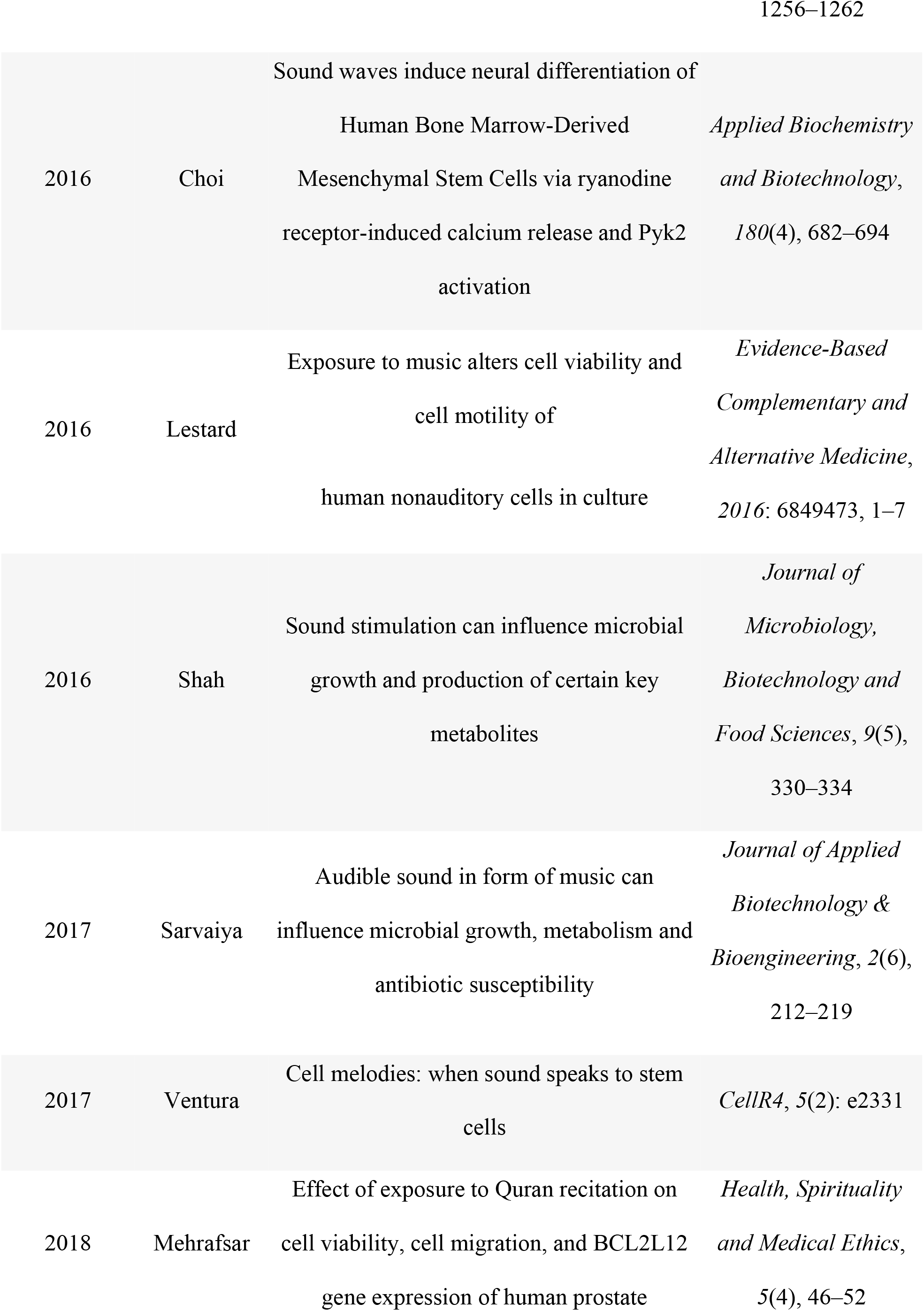
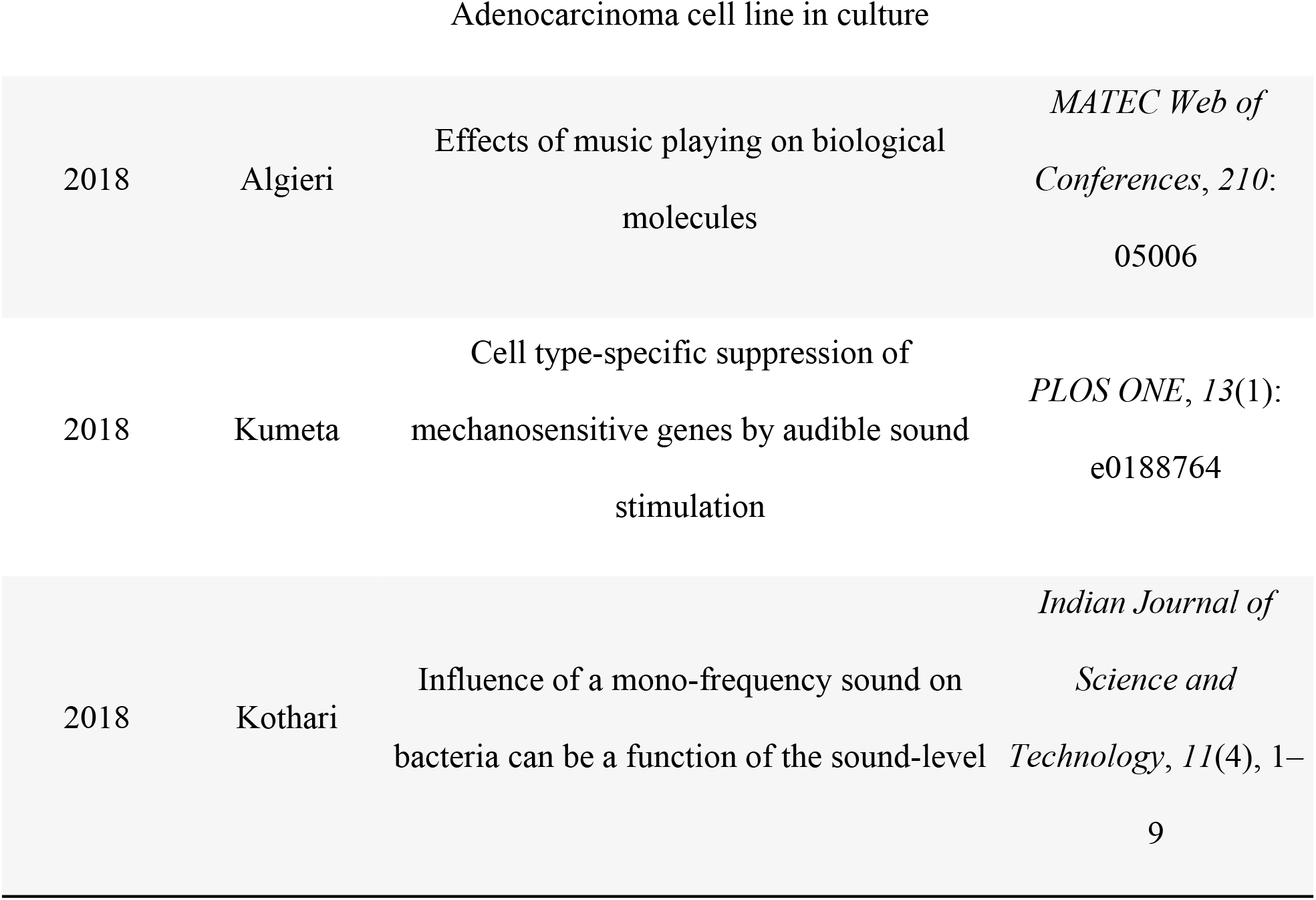
List of included studies.

| Publication<br>year | First author | Title | Source |
| --- | --- | --- | --- |
| 2010 | Shaobin | A pilot study of the effect of audible sound<br>on the growth of *Escherichia coli* | *Colloids and Surfaces*<br>*B: Biointerfaces*,<br>78(2), 367–371 |
| 2012 | Aggio | Sonic vibration affects the metabolism of<br>yeast cells growing in liquid culture: a<br>metabolomic study | *Metabolomics*, 8(4),<br>670–678 |
| 2016 | Mohammed | The effects of acoustic vibration on<br>fibroblast cell migration | *Materials Science and*<br>*Engineering: C*(69), |
| 2016 | Choi | Sound waves induce neural differentiation of Human Bone Marrow-Derived Mesenchymal Stem Cells via ryanodine receptor-induced calcium release and Pyk2 activation | *Applied Biochemistry and Biotechnology*, 180(4), 682–694 |
| 2016 | Lestard | Exposure to music alters cell viability and cell motility of human nonauditory cells in culture | *Evidence-Based Complementary and Alternative Medicine*, 2016: 6849473, 1–7 |
| 2016 | Shah | Sound stimulation can influence microbial growth and production of certain key metabolites | *Journal of Microbiology, Biotechnology and Food Sciences*, 9(5), 330–334 |
| 2017 | Sarvaiya | Audible sound in form of music can influence microbial growth, metabolism and antibiotic susceptibility | *Journal of Applied Biotechnology & Bioengineering*, 2(6), 212–219 |
| 2017 | Ventura | Cell melodies: when sound speaks to stem cells | *CellR4*, 5(2): e2331 |
| 2018 | Mehrafsar | Effect of exposure to Quran recitation on cell viability, cell migration, and BCL2L12 gene expression of human prostate | *Health, Spirituality and Medical Ethics*, 5(4), 46–52 |
| Adenocarcinoma cell line in culture |  |  |  |
| 2018 | Algieri | Effects of music playing on biological molecules | *MATEC Web of Conferences*, 210: 05006 |
| 2018 | Kumeta | Cell type-specific suppression of mechanosensitive genes by audible sound stimulation | *PLOS ONE*, 13(1): e0188764 |
| 2018 | Kothari | Influence of a mono-frequency sound on bacteria can be a function of the sound-level | *Indian Journal of Science and Technology*, 11(4), 1–9 |

## Results

When looking broadly at the selected studies, the sound material used in the literature can be equally divided into the two main sound types identified above:

▪ simple sounds (e.g., sine wave): 6 studies
▪ complex sounds (e.g., music and speech): 6 studies

We will now look more closely at each of these two types of studies, focusing on describing the experimental setup, the sound stimuli used, and the outcomes.

### Simple sound stimulation on cell cultures

In all six studies identified within the category simple sound stimulus, a sine wave with a constant SPL was typically used in various experimental settings. An overview of the findings can be found in Table 2.

**Table 2.**
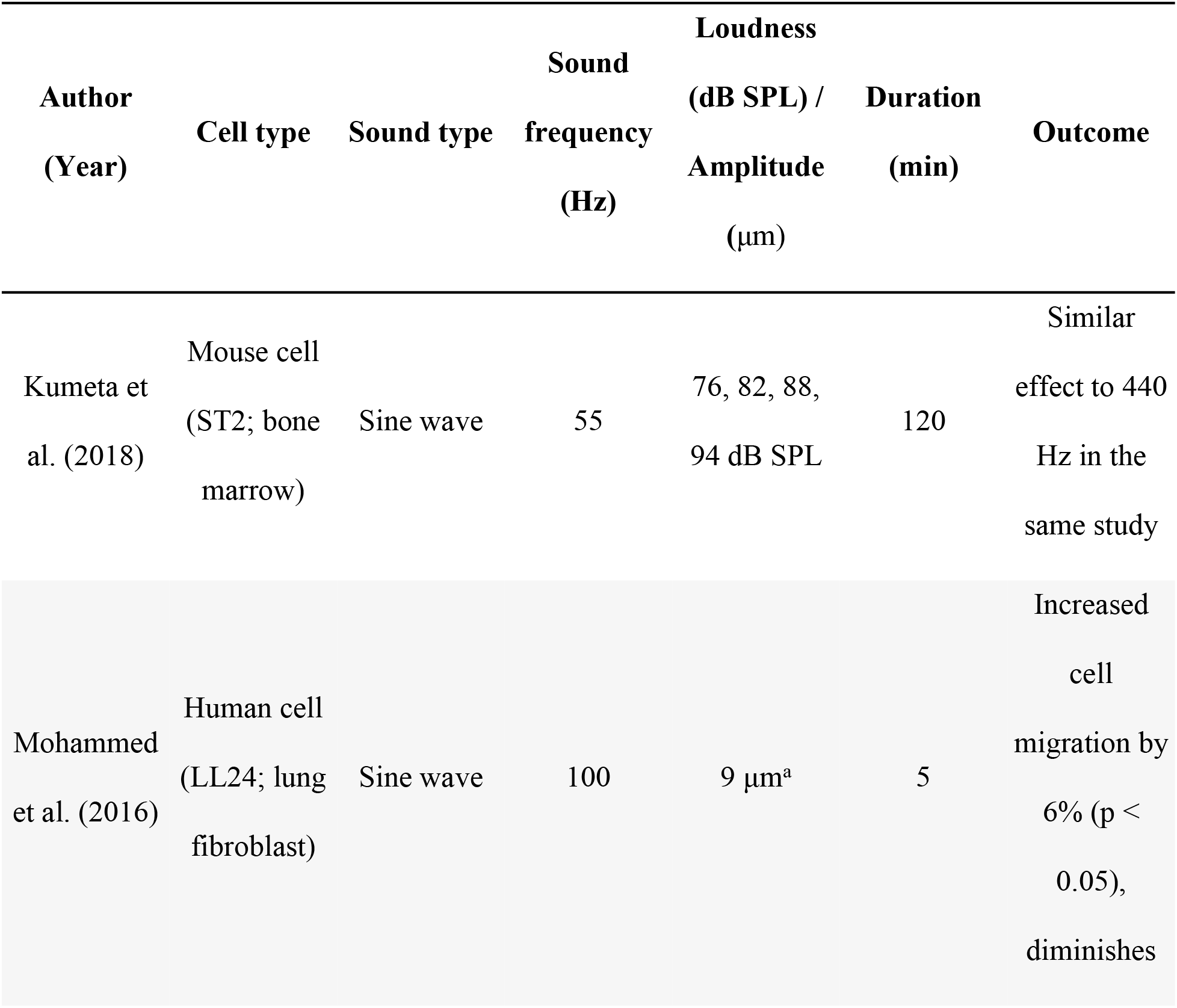
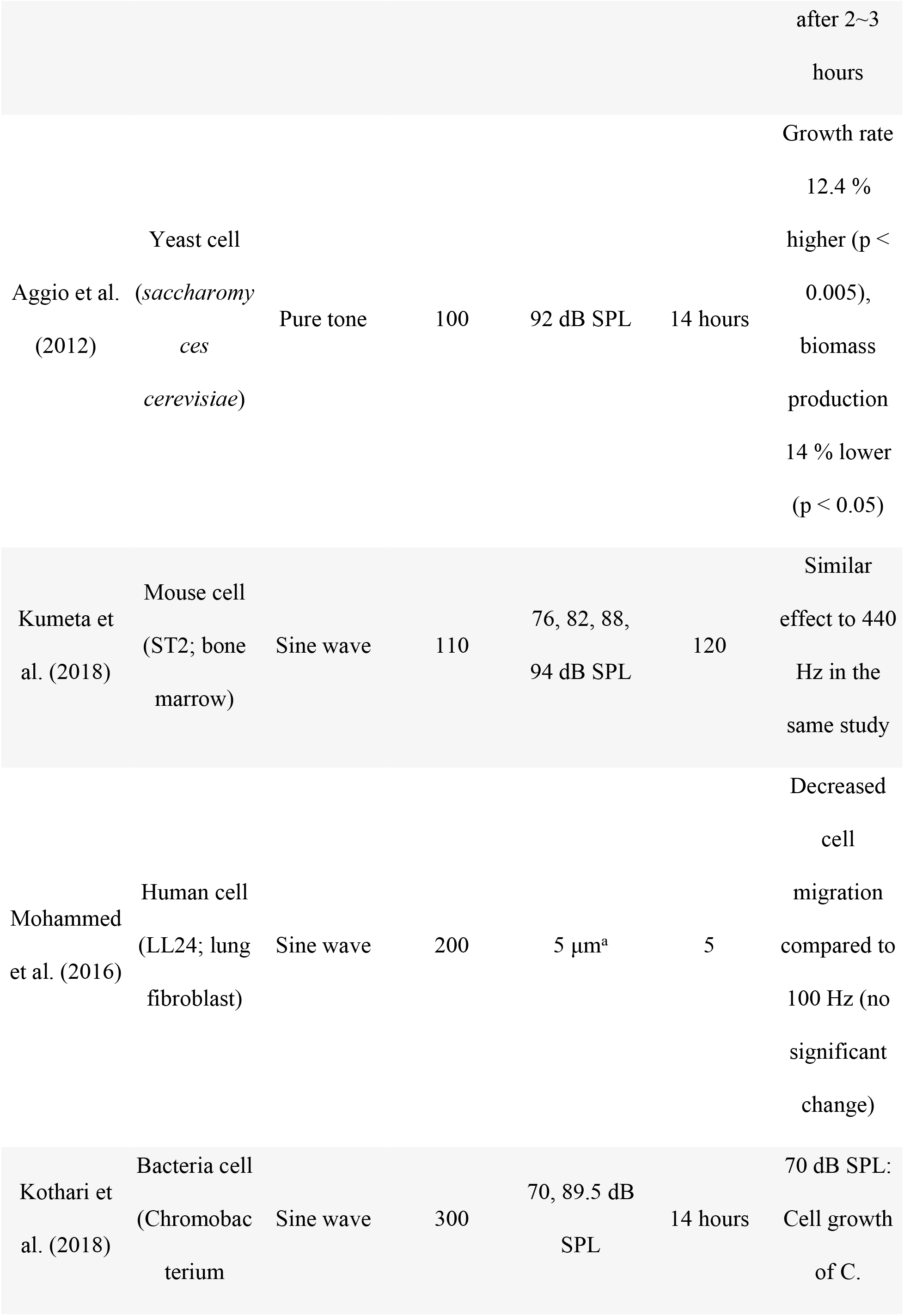
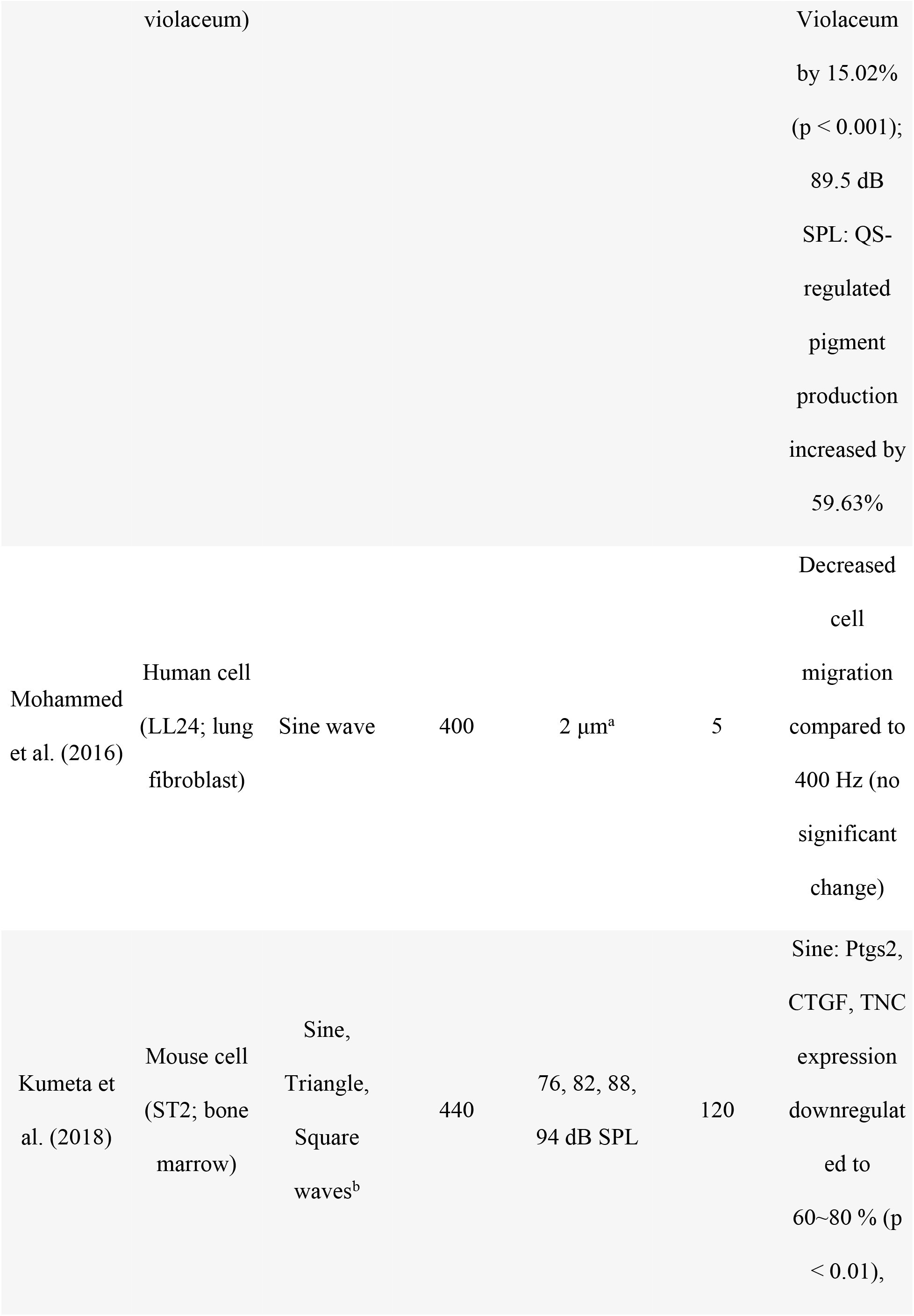
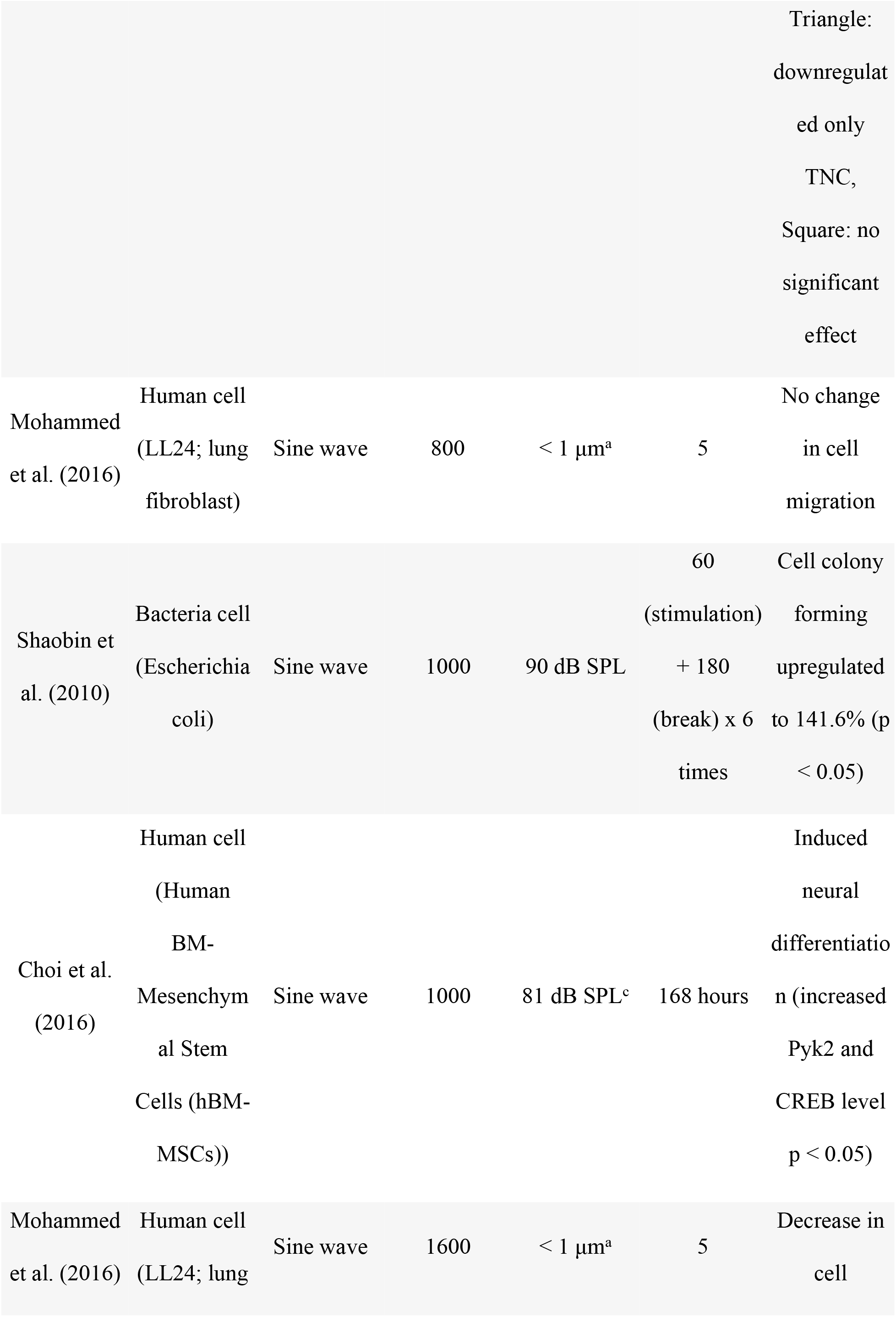
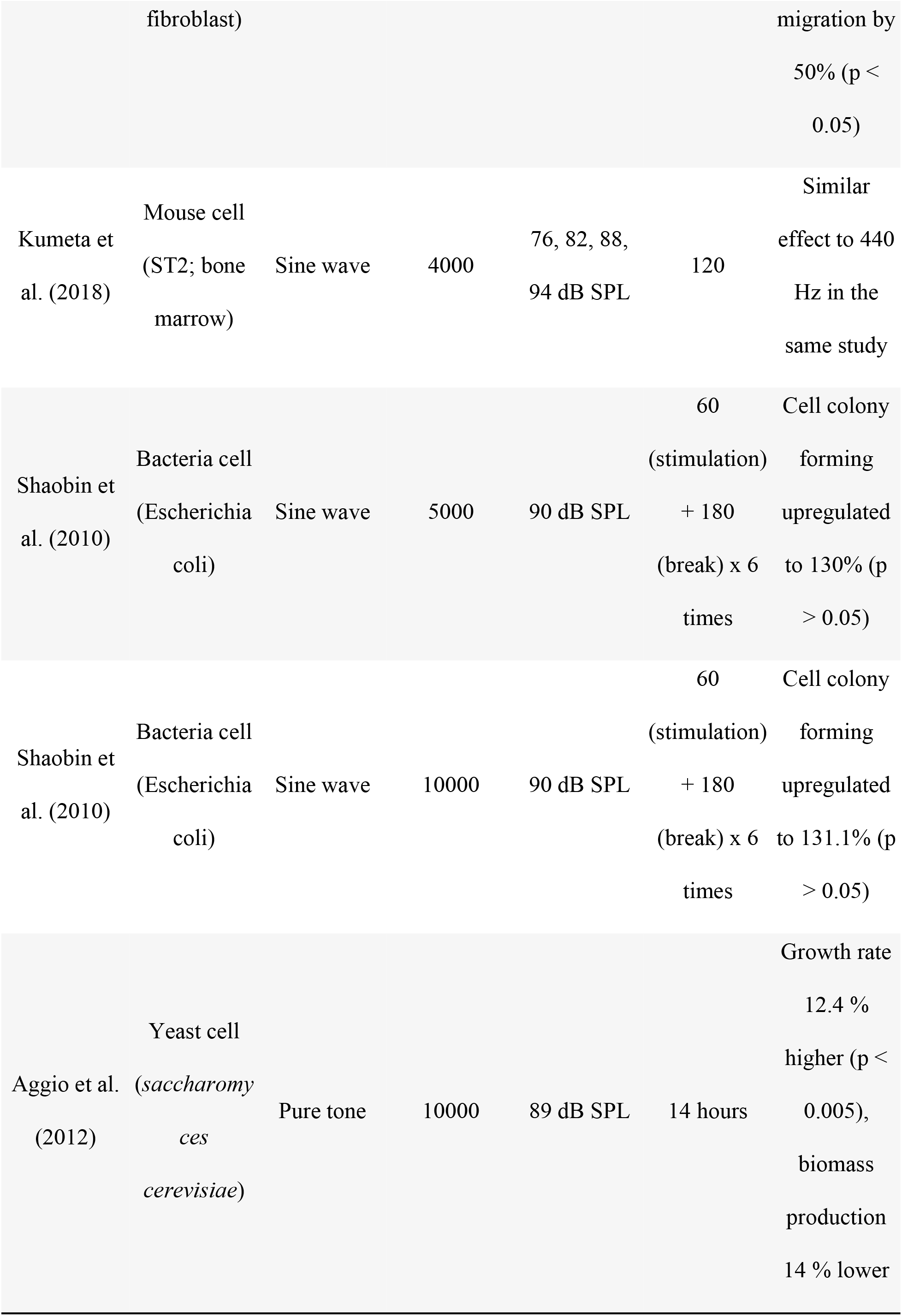
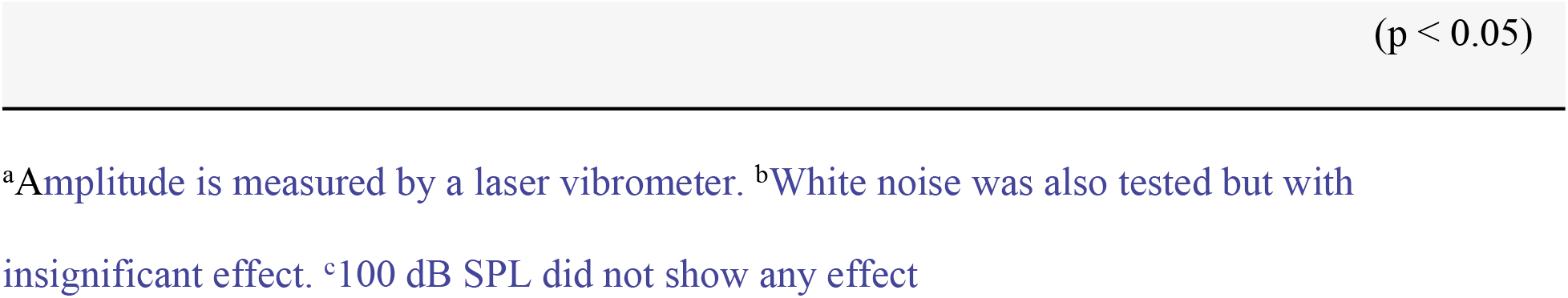
Simple sound stimulus studies (in order of frequency, low to high).

In Shaobin et al. (27), a speaker was placed above the plate inside a box-shaped temperature-controlled (37 °C) environment. The box was acoustically treated with sound absorption installed around the walls. Sine tones with frequencies of 1000, 5000, and 10000 Hz were played separately at 90 dB SPL for 1 hour at a time and repeated 6 times with 3-hour breaks in between. The bacteria cells (*Escherichia coli*) were spread on an agar plate and were therefore not in a liquid-based medium. The cell dish holder was made to rotate using a motor installed outside the chamber. It is not stated whether the control group was placed on the holder while the holder was rotating. Cell colony-forming efficiency was tested, and it was shown that 1000, 5000, and 10000 Hz sine tones upregulated the cell colony-forming by 141.6, 130, and 131.1 % respectively (p < 0.05).

In Aggio et al. (28), a pair of mid-high frequency speakers and a subwoofer was placed around shaker-flasks in a cell incubator. The distance between speakers and flasks was 20 cm. Sine tones of 100 and 10000 Hz were played using computer software at 92 and 89 dBA respectively for 14 hours to a yeast cell culture (*Saccharomyces cerevisiae*). The results showed an effect on the exponential growth rate where it was increased up to 12.4 % (p < 0.005), but after around 10 hours the biomass production was decreased by 14 % (p < 0.05) compared to the control (silence).

In Choi et al. (29), an 8-inch active speaker (NA-818APR, NBN) was placed underneath (10 cm away) the cell culture dish inside a cell incubator. A 1000 Hz sine tone generated by a function generator (FG-7002C, EZ digital) at 81 dB SPL. It was played for 7 days continuously to Human Bone Marrow Mesenchymal Stem Cells (hBM-MSCs). The result showed that the sound treatment induced neural differentiation, and MAP2 and NF-L protein levels significantly increased without altering the cell viability compared to the control group. Phosphorylation levels of Pyk2, ERK, Akt, and CREB were also tested at 10, 45, 90, and 180 minutes post-stimulation. Pyk2 level spiked at 45 minutes and decreased afterwards. The ERK level was increased from 45 minutes and stayed until 180 minutes (p < 0.05). The CREB level was increased from 90 minutes but slightly decreased at 180 minutes (p < 0.05). The Akt level showed no difference. The Pyk2 level was rechecked at a different SPL (100 dB SPL), but did not show any effect, which suggests a possible SPL dependency.

In Mohammed et al. (30), a thin mylar speaker (Ø=45 mm, 0.2 W) was placed directly underneath the cell culture dish (Ø=35 mm). A sine tone with different frequencies (100, 200, 400, 800, 1600 Hz) was generated using an Arduino microcontroller. This induced vibrations in the cell culture dish at a particular displacement amplitude (9, 5, 2, < 1, < 1 μm respectively). The amplitude was measured by using a laser vibrometer (Polytech Ltd.). The dish was resting on the speaker. Different frequencies were applied for 5 minutes each time to two different cell types: human lung fibroblast cells (LL24) and mouse fibroblast cells (L929), placed in separate cell culture dishes. The lowest frequency (100 Hz) was found to increase the migration distance of both cell types by 6 % (p < 0.05), without altering the cell viability. The migration distance decreased at higher frequencies.

In Kothari et al. (31), a speaker (Minix soundbar) was placed 15 cm away next to the cell tubes inside a glass box-shaped chamber (L: 250 x W: 250 x H: 150 mm). The stimuli consisted of sine tones played from the computer software NCH Tone Generator (https://www.nch.com.au/index.html) at 300 Hz and at five different SPLs (70, 76.5, 83, 87.5, 89.5 dB SPL) each for 48 hours. The stimulated cells were bacteria of type *Chromobacterium violaceum*. The noise level was below 40 dB SPL in the control chamber. The result showed an SPL dependency, and the SPL difference of less than 13 dB SPL showed no significant alterations in the growth of cells. The cell growth was upregulated the most at 70 dB SPL, by % (p < 0.001). Quorum Sensing regulated pigment production was affected the most, at 89.5 dB SPL by 59.63 % (p < 0.01).

In Kumeta et al. (8), a 4-inch active speaker (Fostex 6301NB) was placed above (250 mm away) the cell culture dish inside a cell incubator. Different types of sounds (440 Hz sine, triangle and square waves, as well as white noise) were generated at different SPLs (76, 82, 88, 94 dB SPL) using NCH Tone Generator. Each sound was played for 2 hours. The cells in question were of mice (ST2). The background noise inside the incubator was measured to be around 65 dB SPL. The results showed a dependency on the type of waveform used. The sine tones downregulated the gene (Ptgs2, CTGF, TNC) expression level to 60~80 % (p < 0.01). The triangle tones downregulated only one of the genes (TNC), and the square tones and white noise did not show any significant effect compared to the sine tones. Different frequencies (55, 110, 4000 Hz) were also tested for the sine tone condition, but the effect was similar to that of 440 Hz. The SPL of 94 dB SPL showed a significant impact, while tones with other SPL showed little effect. Another important finding in this study was that the temperature changed in the cell culture medium. When sound was played at 94 dB SPL, it caused the temperature in the cell culture medium to increase by 0.4 °C. In this experimental setup, the sound energy was calculated to be reflected by 10.1 dB SPL at the lid of the cell culture dish. Subsequently, the transmitted sound pressure into the liquid-based medium was calculated at approximately 10.4 mPa.

### Complex sound stimulation on cell cultures

Here we will look at studies that have used different types of complex sound stimuli. This includes the use of music (different styles and genres) or speech (human voice) to stimulate the cell cultures in various experimental settings. There were six studies identified in this category. An overview of the findings can be found in Table 3.

**Table 3.**
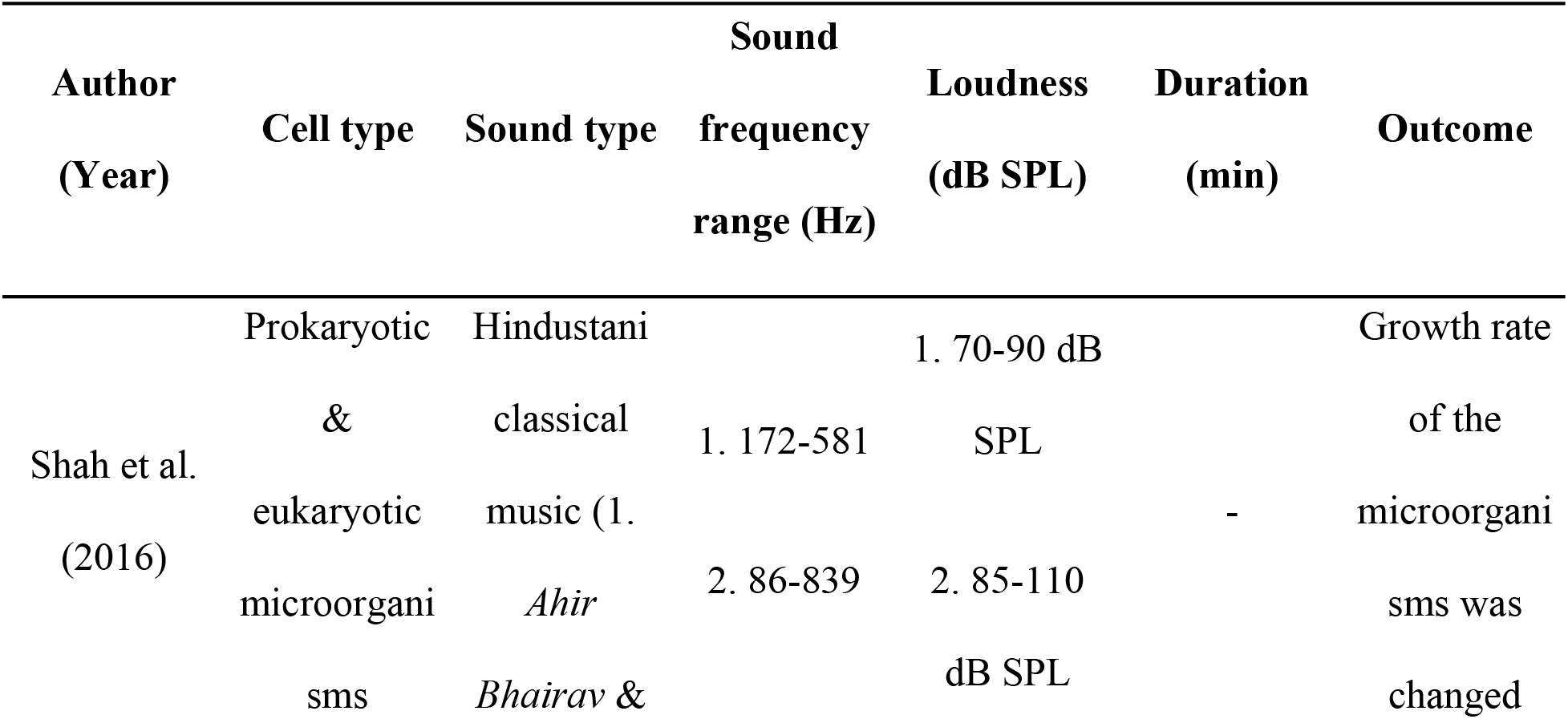
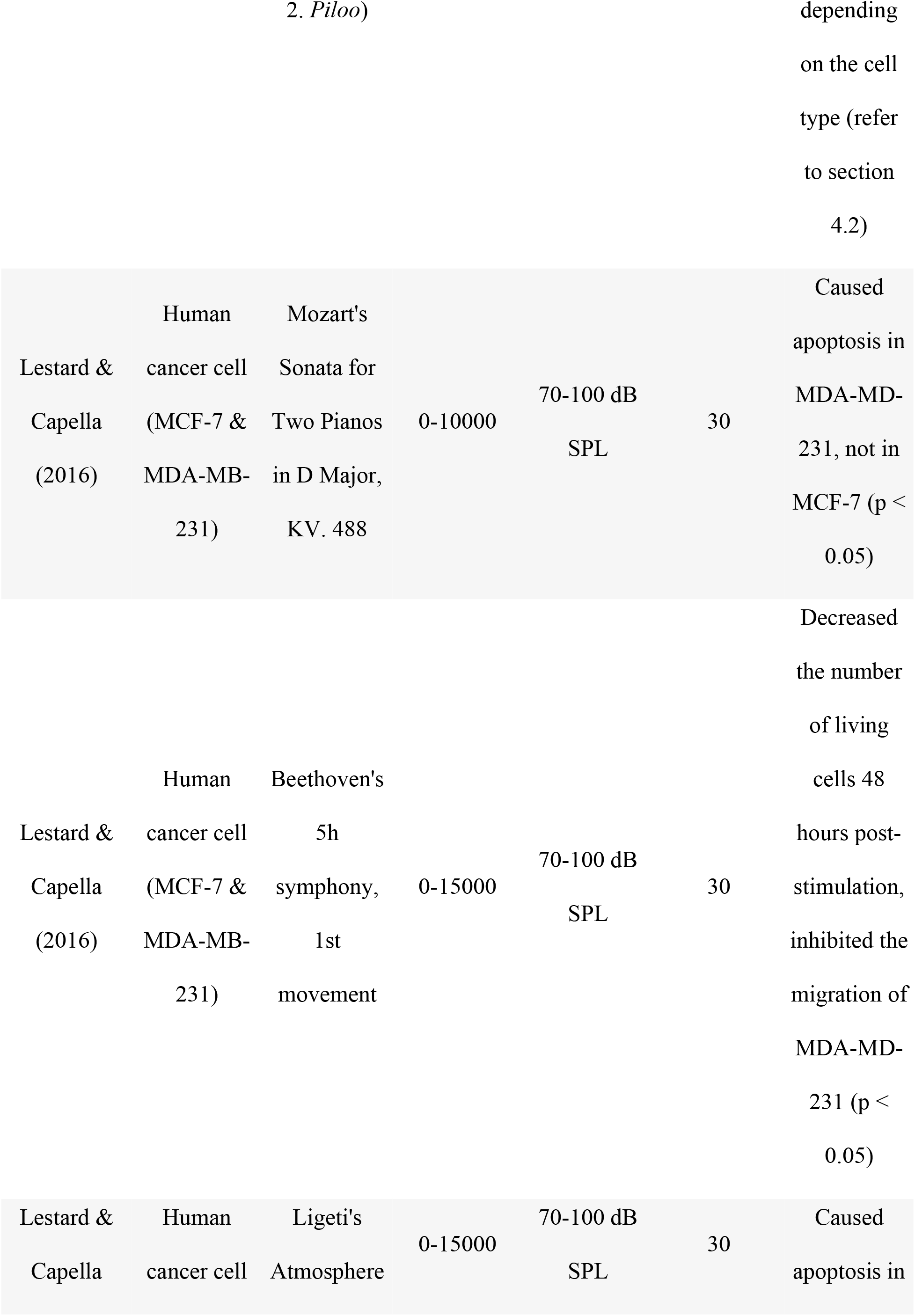
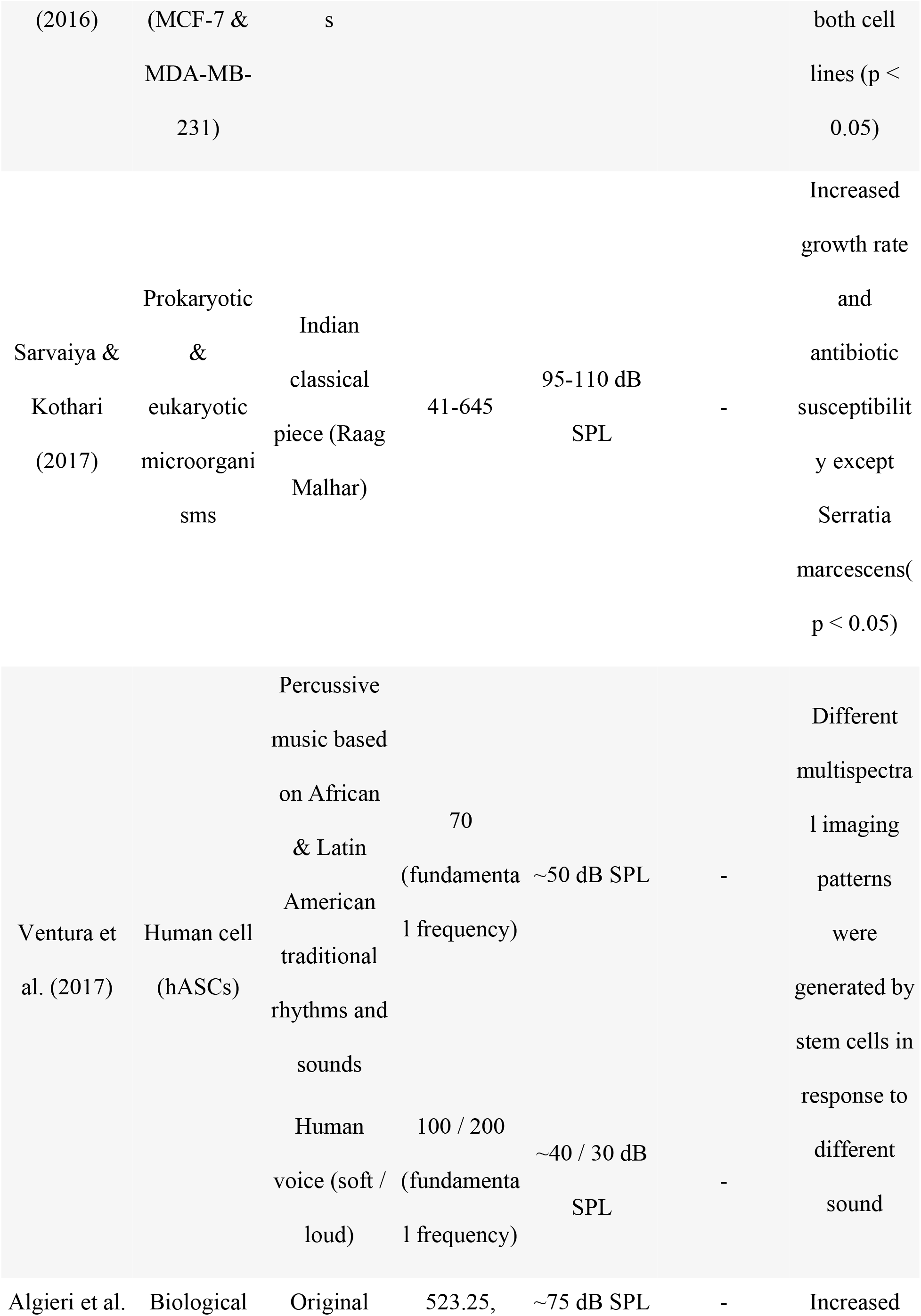
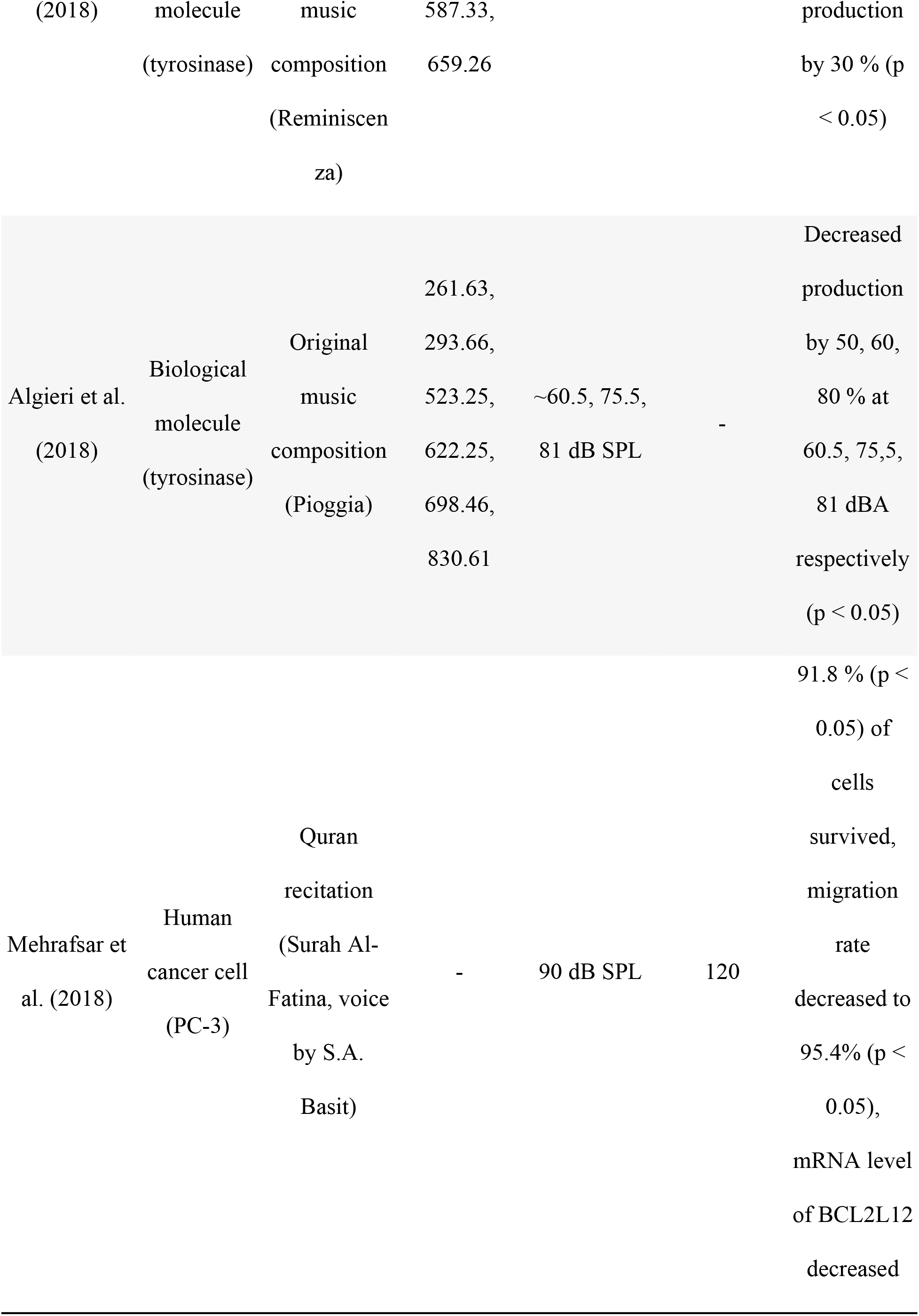
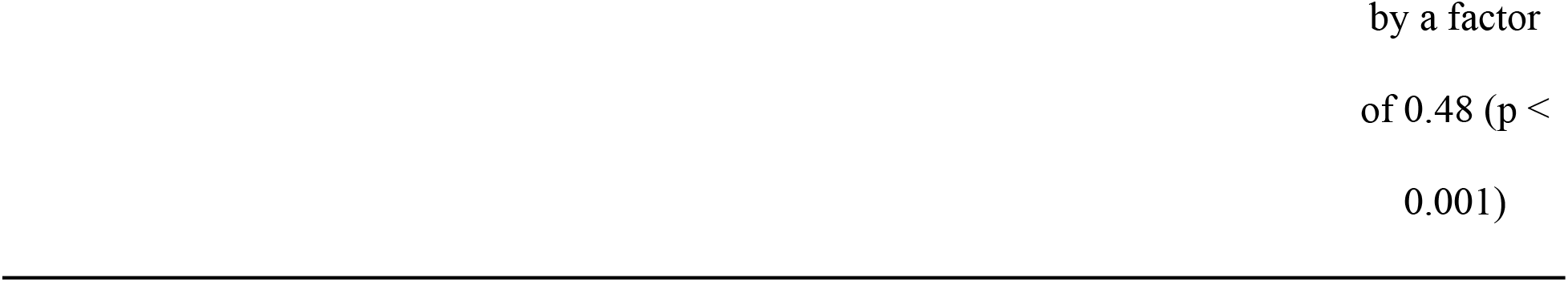
Complex sound stimuli studies (in order of published year).

In Lestard and Capella (32), a 60 W coaxial speaker was installed at the top of the acoustically treated (cork and foam) incubator. The stimuli included the first movement from Mozart’s Sonata for Two Pianos in D Major, KV. 488, the first movement from Beethoven’s 5th symphony, and the first movement from Ligeti’s *Atmospheres.* These excerpts were played at around 70~100 dB SPL for 30 minutes each to human breast cancer cells (MCF-7 and MDA-MB-231). Although it was not reported in the paper, we have found that the frequency range in Mozart’s piece is, approximately, between 0 and 10000 Hz, and 0 and 15000 Hz in both the Beethoven and Ligeti pieces. Mozart’s piece did not alter the viability of MCF-7 but caused cell death (apoptosis) in MDA-MD-231 (p < 0.05). The Beethoven and Ligeti pieces decreased the number of living cells 48 hours post-stimulation and inhibited the migration of MDA-MD-231 (p < 0.05). The Ligeti piece caused significant cell apoptosis in MCF-7 (p < 0.05).

In Shah et al. (33), a speaker was placed next to (15 cm away) the cell tubes inside a glass chamber (225 × 225 × 125 cm). An SPL meter (ACD Machine Control Company) was positioned 15 cm away from the speaker. Two Hindustani classical pieces *Ahir Bhairav* and *Piloo* were played at 70~90 dB SPL and 85~110 dB SPL, respectively, to six different microorganisms (prokaryotic and eukaryotic). The recordings were from a CD album named *Call of the Valley* (Saregama Indian Ltd.). The fundamental frequency ranges of the two pieces were 172~581 Hz (*Ahir Bhairav*) and 86~839 Hz (*Piloo*). After playing *Ahir Bhairav*, the growth rate of *Serratia marcescens* (−4.32 %, p < 0.01), *Chromobacterium violaceum* (4 %, p < 0.05), *Xanthomonas campestris* (6.66 %, p < 0.05), *Saccharomyces cerevisiae* (8.94 %, p < 0.01), *Lactobacillus plantarum* (6.14 %, p < 0.01), and *Bacillus parabrevis* (18.75 %, p < 0.01) were changed compared to the control. After playing *Piloo*, the growth rate of *Serratia marcescens* (−7.46 %, p < 0.01), *Chromobacterium violaceum* (−24.19 %, p < 0.01), *Xanthomonas campestris* (9.32 %, p < 0.05), *Saccharomyces cerevisiae* (4.68 %, p < 0.01), *Lactobacillus plantarum* (3.41 %, p < 0.01), and *Bacillus parabrevis* (18 %, p < 0.05) were changed compared to the control.

In Sarvaiya and Kothari (34), the experimental setup was the same as in Shah et al. (2016). A Hindustani classical piece (Raag Malhar) was played at around 95~110 dB SPL to various microorganisms (prokaryotic & eukaryotic). The fundamental frequency range of the music was between 41 and 645 Hz. All tested cells showed an increase in growth rate, and in antibiotic susceptibility, except *Serratia marcescens* (p < 0.05). The alcohol tolerance of *Saccharomyces cerevisiae* was significantly enhanced by 33.69 % (p < 0.001) after 24 hours of incubation when the concentration of ethanol was 5 %v/v, 132.94 % (p < 0.001) after 24 hours of incubation when the concentration of ethanol was 10 %v/v, 27.92 % (p < 0.01) after 48 hours of incubation when the concentration of ethanol was 12.5 %v/v, and 1200 % (p < 0.001) after 120 hours of incubation when the concentration of ethanol was 15 %v/v.

In Ventura et al. (18), the cell culture was placed two meters in front of a live performance. Percussive music based on traditional rhythms and sounds from Africa and Latin America and an actor’s voice were played for 45 minutes and 30 minutes, respectively to Human Adipose Stem Cells (hASCs). The tempo of the percussive music was taken from the performer’s own heartbeat, and the actor’s voice was performed with two different intensities (soft and loud). The fundamental frequency for the percussive music was around 70 Hz at ~50 dB SPL, and for the actor’s voice was around 100 Hz at ~40 dB SPL (soft voice) and 200 Hz at ~30 dB SPL (loud voice). The sound analysis was done through Praat, a software package (www.fon.hum.uva.nl/praat/) developed for speech analysis. The cell culture was observed in real-time under the Multi Spectral Imaging (MSI) system (Nikon Eclipse TS 100). The MSI analysis showed spectral changes of cell imaging with specific vibrational signatures in relation to the stimuli.

In Algieri et al. (7), a set of 3 W internal speakers (LENOVO E93z) was placed underneath (10 cm away) the reactor vessel on a magnetic plate. The SPL measurement was done with a FUSION −1 dB meter and analyzed using dB Trait. Two distinctively composed pieces of music, *Reminiscenza* (constant rhythm and melody) and *Pioggia* (irregular rhythm with more variable devices such as irregular tempo changes, and melody) were played to biological molecules (tyrosinase). These were placed in a reactor vessel with a stirring mechanism and a constant temperature of 30 °C. In *Reminiscenza*, only C5 (523.25 Hz), D5 (587.33 Hz), and E5 (659.26 Hz) musical notes were used and played at ~75 dBA. In *Pioggia*, C4 (261.63 Hz), D4 (293.66 Hz), C5 (523.25 Hz), D#5/Eb5 (622.25 Hz), F5 (698.46 Hz), and G#5/Ab5 (830.61 Hz) were used, and played at different SPL (60.5, 75.5, and 81 dBA). The 1-DOPA production rate was tested. Playing *Reminiscenza* resulted in a significant increase in production, by 30 % more compared to the control group. Playing *Pioggia* resulted in decreased production by 50 % at 60.5 dBA, 60 % at 75.5 dBA, and 80 % at 81 dBA (p < 0.05).

In Mehrafsar et al. (35), four speakers were placed around the cell culture plate. A recording of Quran recitation (*Surah Al-Fatina,* voice by *Sheikh Abdul Basit*) was played at 90 dB SPL for 2 hours to the human cancer cell line (PC-3; Human Prostate Adenocarcinoma Cells). The cell viability, migration, and BCL2L12 gene expression level were tested. After the sound stimulation, 91.8 % (p < 0.05) of PC-3 cells survived, the migration level decreased to 95.4 % (p < 0.05), and mRNA level of BCL2L12 decreased by a factor of 0.48 (p < 0.001).

## Discussion

As we have seen from the various studies above, many different types of both simple and complex sound types, with an extensive range of frequencies and SPLs have been tested. The results reported for both simple and complex sound stimulation studies are promising, either they can enhance cell migration, proliferation, colony formation, and differentiation ability or decrease cell migration and cause apoptosis of cancer cells. However, it is challenging to find a systematic pattern and to compare the results because the studies use widely different sound materials, experimental setups, sound measurement methods, and observation strategies. Below, we synthesize and discuss the results while considering the practical matters (e.g., experimental setups and measurement methods) and also the physics of sound.

### Experimental Setup

Many of the reviewed studies were based on a speaker position above, underneath or next to the cell cultures. The distances varied, ranging from ~0 (directly underneath the dish) to 200 cm. All three placements resulted in cellular responses without any apparent difference. It would be of value to perform a systematic experiment of how different positioning of the sound source (the speaker) results in various types of mechanical force in the cell medium, for example whether or not a speaker is coupled (in direct contact) with the cell dish or plate. In terms of the acoustic treatment of the experimental environment, this is sometimes not clearly stated. Based on its physical properties, a sound stimulus may interact with the testing environment. Ideally, sound-related experiments should be performed in an acoustically controlled environment. The ideal solution might be to use an anechoic chamber, which is designed to imitate the free field conditions (i.e., no reflected sound waves), and which would allow no or minimum external sound to enter the chamber (36). This would improve the measurements of the dependent variable and ultimately help reach conclusions about how different variables are correlated (37). Also, for practical reasons, speakers should ideally be placed inside an acoustically treated chamber in order not to damage the researcher’s hearing in the lab. For example, a continuous sine tone at a loudness level of 70 dB SPL or higher can cause temporary or even permanent hearing damage (21). In most cases, an anechoic chamber would not be feasible considering the cost and effort. However, it would still be useful to try to reduce the acoustic impact of the environment by using some kind of damping element. It would also be essential to measure how the sound stimulus behaves in the environment. This could be used to address potential obstacles during experiments, for example, amplitude distortion at specific frequencies, standing waves, and phase cancellation.

### Sound features

We will look at four basic sound features here: frequency (Hz), sound pressure level (dB SPL), and waveform (simple & complex).

#### Frequencies

In the studies where single-frequency sound stimuli were used, single frequencies selected in the range of 55~10000 Hz were explored on bacteria cells (*Escherichia coli*, *Chromobacterium violaceum, Serratia marcescens*), mouse cells (ST2), and human cells (hBM-MSCs, LL24). In the studies where the more complex sound stimuli were used, musical pieces or human voice in the frequency range between 0 and 15000 Hz were explored on biological molecule (tyrosinase), microorganisms (various prokaryotic & eukaryotic), bacteria cells (*Escherichia coli*), human cancer cells (MCF-7, MDA-MD-231), and human cells (hASCs). To summarize, bacteria and yeast cell studies showed an effect of sound across a wide range of frequencies: cell colony-forming and growth rate increase using 100, 300, 1000, 5000, and 10000 Hz (28,29,32). Mammal cells studies showed a little more selective and frequency-dependent effect. For example, a lower frequency (100 Hz) induced cell migration increase in human cells, but higher frequencies (above 1600 Hz) induced the opposite effect on cell migration (30). Also, stimuli of 440 Hz induced gene (Ptgs2, CTGF, TNC) expression increase in mouse cells (ST2), while 55, 110, and 4000 Hz showed a smaller but similar effect (8).

For the complex sound stimulation studies, there are too little reports in the literature, but the results are intriguing. The tested music and human voice varied widely: Western classical music such as Mozart, Beethoven, and Ligeti (32), Hindustani classical music such as *Ahir Bhairav, Piloo,* and *Raag Malhar* (33,34), an African and Latin American percussive music and a voice of an actor (18), a contemporary composition such as *Reminiscenza* and *Pioggia* (7), and Quran recitation such as *Surah Al-Fatina* (35). The most present frequencies of the music and voices tested stayed between 40 and 840 Hz. However, it is most likely that other frequency content, especially overtones, varies between different types of music or speech style due to distinctive *timbre* differences between instruments and human voices. Since complex sounds typically consist of many frequencies, it is difficult to know if a particular frequency had an impact on the outcome. However, a notable reported outcome is where two musical compositions of contrasting characteristics in terms of melody and rhythm (regular/constant vs irregular/spontaneous) showed contrasting results in 1-DOPA production rate in *tyrosinase*. Music with regular characteristics (constant harmonic and rhythmic structure) using 523.25 Hz (musical note: C5), 587.33 Hz (D5), and 659.26 Hz (E5) resulted in 1-DOPA production increase. Music with more irregular characteristics (disharmonic and irregular rhythm and tempo structure), on the other hand, using 261.63 Hz (C4), 293.66 Hz (D4), 523.25 Hz (C5), 622.25 Hz (D#5/Eb5), 698.46 Hz (F5), and 830.61 Hz (G#5/Ab5), resulted in 1-DOPA production decrease (7). This indicates a possible role of the degree of pitch (frequency), also possibly of harmonic versus disharmonic frequency content, and temporal regularity in the produced outcome.

#### Sound pressure level

In the investigated studies, the sound pressure level range used varied extensively, from 40 to 110 dB SPL. The majority of the studies used the range between 75 and 95 dB SPL, which is as loud as a vacuum cleaner (3 meters away) and a subway train (6 meters away), respectively. The range centered around 90 dB SPL, which is about 632 mPa. This is above the loudness level where sustained exposure can cause hearing damage. However, the pressure is small compared to the periodic blood flow pressure, for example, which can typically be around 120/80 mmHg (approximately 10~16 kPa) (38). The transmitted sound energy in the cell medium from air is estimated to be even smaller: approximately 29.67 mPa (when the temperature is 20 °C, *R*≈0.9988, transmitted sound energy from the air to water is 1−*R*≈0.0022 when there is no absorption). This is a small mechanical stress, but the cells have mechanisms that can sense such small forces and changes, for example, α-catenin (8,39).

It should be noted that some of the SPLs specified in the literature are incomparable. This is because some of the reports were unclear about how the actual sound output was measured. Additionally, some of the commercially available SPL measurement devices typically come with filters that imitate the human hearing characteristics (less sensitive at very low and high frequencies within the audible frequency range), such as an A-weighting filter. There are other weightings, as illustrated in Fig 2 (40), designed for different applications. This is a small piece of information that may not be considered necessary, but it should be stated in the report, as a specific weighting filter readout can undermine a particular frequency range as shown.

**Fig 2.**
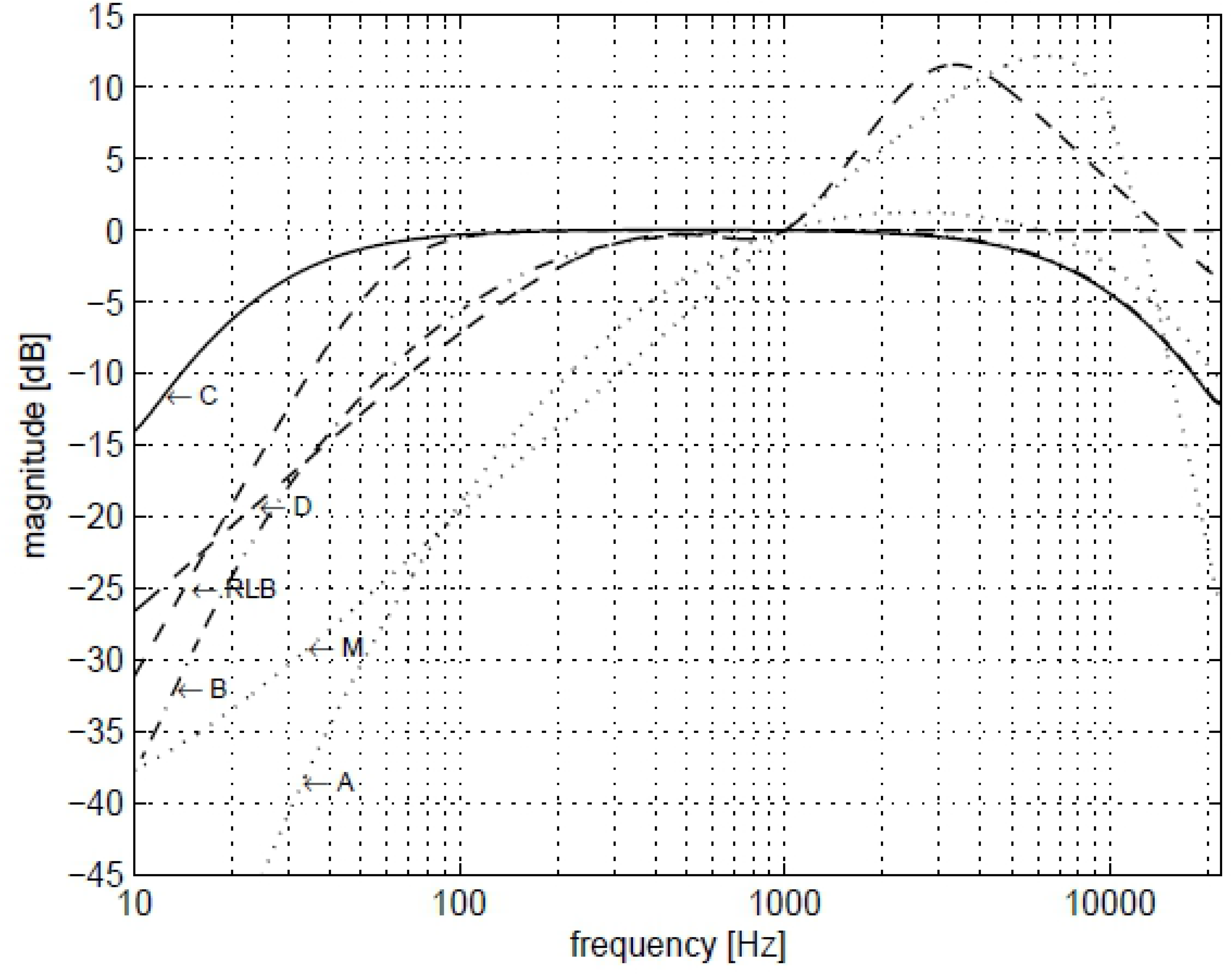
The frequency responses of A, B, C, D, M and RLB weighting filters (40).

#### Waveform

As pointed out earlier, the majority of the simple sound studies employed sine tones. There was only one study that compared different types of simple waveforms (sine, triangle, square) and white noise. Interestingly, the different waveforms yielded contrasting outcomes in cellular responses (8). The underlying mechanism is not well understood, but an analysis of the waveforms can be made: sine waves are continuous, whereas triangle and square waves are discontinuous. Also, sine waves do not have any harmonic frequencies (overtones), only its fundamental frequency. Triangle and square waves, on the other hand, contain odd harmonics above its fundamental frequency. The higher harmonics of the triangle wave get weaker faster than those of a square wave, as shown in Fig 3. Due to its wave shape, the amount of time that the square wave stays in its maximum amplitude is longer than other waveforms. This is a reason why square waves are perceived louder than sine waves. When the amplitude and the frequency are the same, there is more sound energy in square waves and stronger harmonics than in sine and triangle waveforms. Although there is a need for further investigation, according to the differences we explore, a possible explanation of different outcomes between sine, triangle, square, and white noise could be the strength of harmonics present in the sound material.

**Fig 3.**
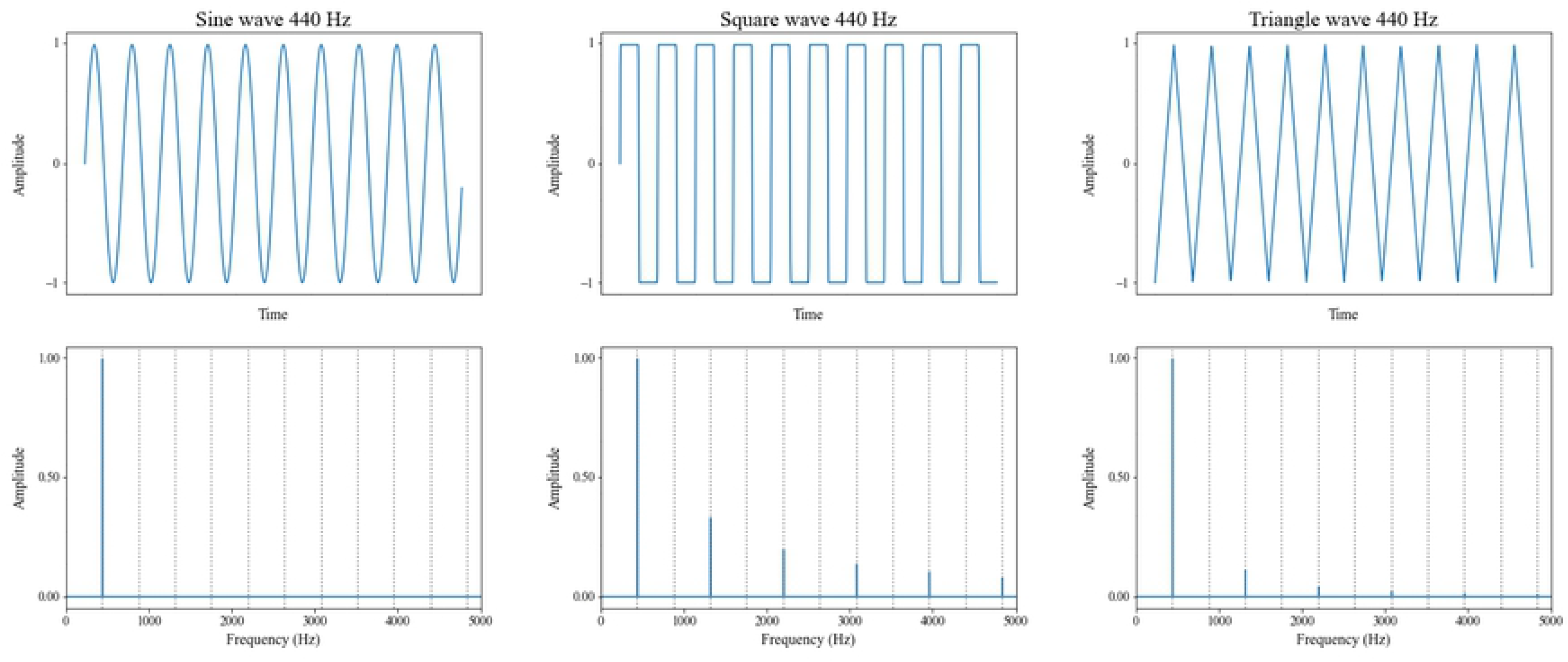
Comparison of sine, triangle, and square waves with a fundamental frequency of 440 Hz.

### Rhythms

Rhythm is one of the fundamental properties of music and is also an essential feature of biological life. In the investigated simple sound studies, there were no rhythmic features in the sound stimuli. On the other hand, all the complex sound studies had an aspect of rhythm. The rhythms varied widely: Western classical music, Hindustani classical music, and African and Latin-American traditional music. For illustration, Mozart’s Sonata for Two Pianos in D Major, KV. 488 shows more regular and steady rhythmic patterns than Ligeti’s *Atmospheres* (Fig 4a & Fig 4b). There was only one study in which the aspect of rhythm was deliberately tested and reported, and where regular/constant and irregular/inconstant rhythms were compared in terms of cellular responses (7). The results showed that the musical piece with regular rhythm produced an increase in the production of 1-DOPA compared to the control group, whereas the piece with irregular rhythms resulted in a decrease.

**Fig 4a.**
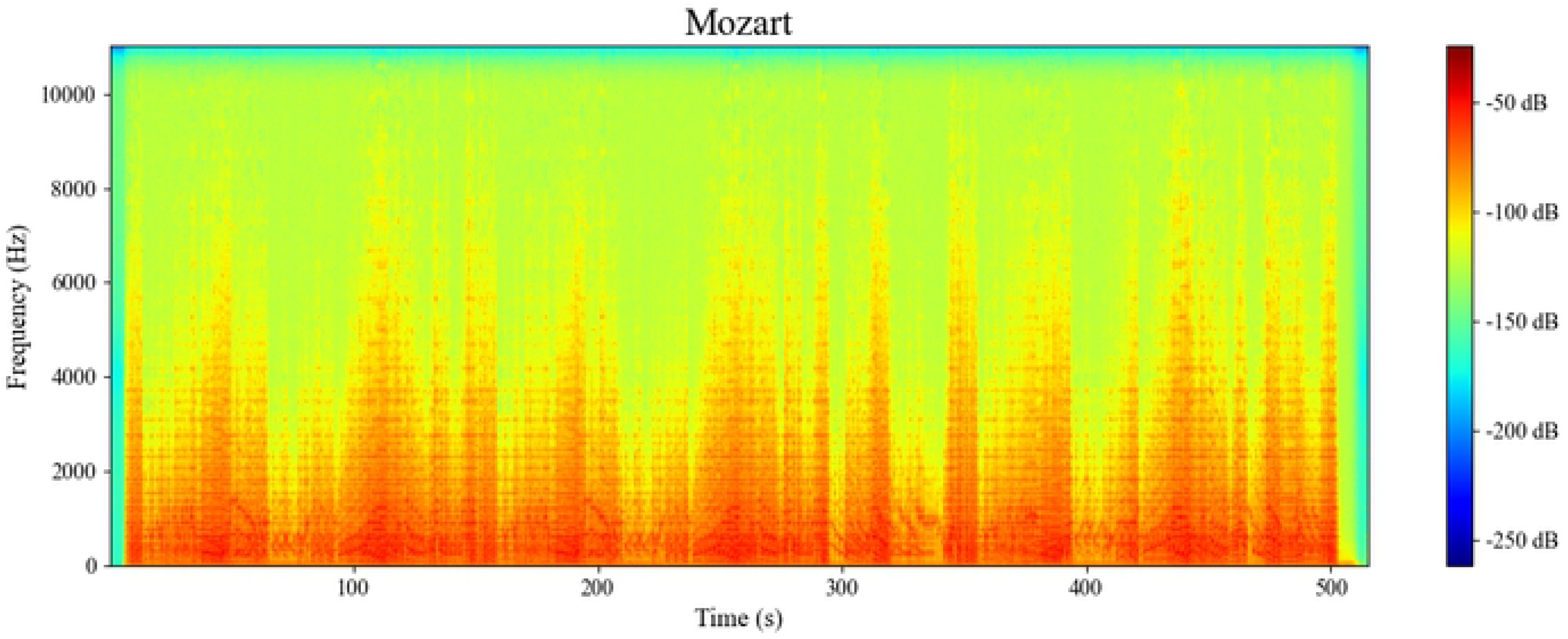
Spectrogram of the first movement of Mozart’s Sonata for Two Pianos in D Major, KV.488.

**Fig 4b.**
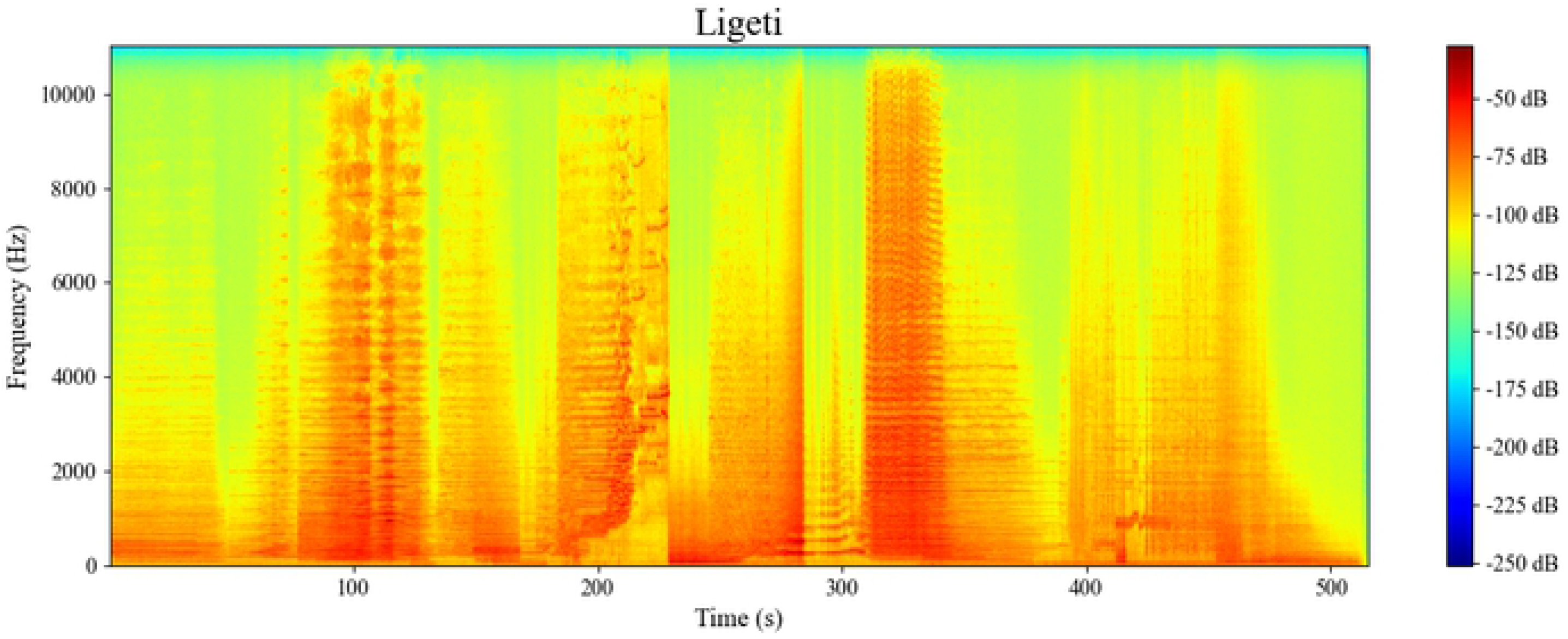
Spectrogram of Ligeti’s Atmospheres.

The “synchronization of multiple rhythms” is vital in living organisms (41), and rhythmicity is one parameter that is underexplored in the literature. The human body is full of different rhythms, such as heartbeats, which are paced by the so-called sinus node and respiratory rhythms that are controlled by the bilateral dorsal respiratory groups. If certain rhythms in the human body become irregular, it can be an indication of more severe health problems, for example, arrhythmia (42) or abnormal respirations (43). Together, rhythmicity is a possible new avenue of sound stimulation of cells and is of value to be investigated further.

## Conclusion

As we have seen from this literature review, there appear to be promising effects of using audible sound on microorganisms and mammalian cells. The correlations are complex, and the exact cause of cell responses to audible sound stimulation is still largely underexplored. We have also found that there are considerable differences in experiment setups and sound features when it comes to the setups used for cellular stimulation. In many cases, it is also unclear exactly how the stimulation happened due to an uncontrolled (from a sonic point of view) environment. In future research, we, therefore, see the need for:

- **Controlled sonic environment**: An acoustically treated environment would make it easier to control the sound stimuli. This would reduce the level of external noise and avoid reinforcement or cancellation of particular frequencies. What is being applied to the cells should only be the sound emitted directly from a speaker. The ideal would be to use an anechoic chamber, but even installing suitable dampening acoustic materials would improve the experimental environment.
- **Measurement method**: Due to the possible (sonic) influence of the environment as mentioned above, it is necessary to measure and analyze the environment with appropriate equipment (e.g., a flat frequency response calibration microphone). This would serve as a control, and an SPL measurement method should be used to know what the cells are stimulated with.
- **Sound stimuli**: A more comprehensive range of sound stimuli should be tested. All of the simple sound stimuli used in the literature so far have been of a continuous nature. Considering the complexity of the extracellular matrix (environment) that the cells live in, learning the influence of pulsating or rhythmic stimuli on cells could also be interesting. The control of the sound factors/parameters that are varied should be improved when using more complex sound stimuli.

Many other possible non-sound related factors can alter the functionality of cells under examination (44). Therefore, as to recommendations for future research in this multidimensional and uprising field, it is crucial to have the sound stimuli well under control and elucidate the practical issues. High priority should be given to investigate the optimal acoustic environment for cell stimulation that takes cell viability (e.g., temperature) into consideration. Focus should also be put on standardizing methods to measure and analyze the characteristics of the experimental sound output and the background noise. Finally, we suggest replication studies of the findings from the literature under the above-mentioned controlled conditions.

